# Quantifying Differences in Neural Population Activity With Shape Metrics

**DOI:** 10.1101/2025.01.10.632411

**Authors:** Joao Barbosa, Amin Nejatbakhsh, Lyndon Duong, Sarah E. Harvey, Scott L. Brincat, Markus Siegel, Earl K. Miller, Alex H. Williams

## Abstract

Quantifying differences across species and individuals is fundamental to many fields of biology. However, it remains challenging to draw detailed functional comparisons between large populations of interacting neurons. Here, we introduce a general framework for comparing neural population activity in terms of *shape distances*. This approach defines similarity in terms of explicit geometric transformations, which can be flexibly specified to obtain different measures of population-level neural similarity. Moreover, differences between systems are defined by a distance that is symmetric and satisfies the triangle inequality, enabling downstream analyses such as clustering and nearest-neighbor regression. We demonstrate this approach on datasets spanning multiple behavioral tasks (navigation, passive viewing of images, and decision making) and species (mice and non-human primates), highlighting its potential to measure functional variability across subjects and brain regions, as well as its ability to relate neural geometry to animal behavior.

## 1 Introduction

Measuring individual differences within and across species is fundamental to many areas of biology (Nelson 1970; Wagner 1989). Such comparisons have been highly successful in certain areas of neuroscience, such as in surveys of gross-level neuroanatomy across species (Butler and Hodos 2005) and in studies of small and self-contained central pattern generator circuits (Marder and Goaillard 2006). However, it remains challenging to quantitatively compare the dynamics of larger neural systems, such as those within mammalian brains.

Over the past couple of decades, a multitude of methods have been proposed to make these comparisons at the level of neural population dynamics. Pioneering work developed Representational Similarity Analysis (RSA; Kriegeskorte et al. 2008), drawing inspiration from existing concepts in cognitive psychology (Edelman 1998). Underscoring the complexity of this topic, several variants of RSA were proposed by subsequent work (see Kriegeskorte and Diedrichsen 2019, for a review), while others pursued alternative methods, such as: Canonical Correlations Analysis (CCA; Raghu et al. 2017; Gallego et al. 2020) and its generalizations (reviewed in Zhuang et al. 2020), Procrustes alignment (Degenhart et al. 2020), and hyperalignment (Haxby et al. 2011; Haxby et al. 2020). Studies of neural computation in artificial deep networks have utilized similar techniques, notably Centered Kernel Alignment (CKA; Kornblith et al. 2019), which was later found to be closely related to RSA (Williams 2024). Other works have used linear regressions to compare encoding geometries across brain regions (e.g. Bernardi et al. 2020) and to quantify similarity between artificial and biological network representations (e.g. Conwell et al. 2022). A practitioner who wishes to compare the representational geometries of multiple neural circuits is therefore faced with a rich, but complex and interrelated, ecosystem of quantitative methods. It is often unclear how these various methods relate to each other, or whether they are each special cases of a unifying framework.

Scaling up these analyses to compare large collections of neural systems remains a challenge. For example, the International Brain Lab (IBL) released a dataset consisting of 121 recording sessions in 78 mice performing a visual discrimination task (International Brain Laboratory et al. 2024). The methodologies summarized above can quantify similarity between any pair of sessions (e.g. by computing the average canonical correlation coefficient). However, these similarity scores do not provide many direct insights in and of themselves. Indeed, the 121 sessions in the IBL dataset yield 7260 similarity scores between pairs of recording sessions. How should researchers make sense of these score values? One possibility is to show that these scores are, in aggregate, more similar to each other than a null control group (constructed, e.g., by shuffling data). But these simple approaches do not chart a path towards answering more detailed questions. For instance, it would be useful to test whether these scores indicate the presence of clusters—i.e., small groups of subjects with similar neural activity to each other. Additionally, it would be useful to use these scores to make predictions—e.g., to infer the behavioral profile of an individual subject from that subject’s neural similarity to others.

Here, we introduce a principled approach to geometrically compare activity measurements among multiple neural populations (e.g. many different brain regions or animal subjects), which is broadly referred to as *shape distance*. Through analyses of simulated and experimental datasets we highlight two major advantages of this approach. First, the shape distance between a pair of systems is quantified in terms of a transparently specified class of neural alignments (e.g. rotations + reflections, linear, or nonlinear transformations), thereby providing a flexible and interpretable family of measures. Second, each distance within the shape metrics framework satisfies a couple of key mathematical properties—symmetry and triangle inequality—which allow rigorous clustering and regression analyses across large numbers of recordings.

The mathematical foundations of the shape metrics framework were laid out in a series of recent conference papers (Williams et al. 2021; Pospisil et al. 2023; Harvey et al. 2024; Khosla and Williams 2024). The present manuscript substantially expands upon these prior works by demonstrating the methodology on practical applications in neuroscience, spanning comparisons across brain regions and individuals, in rodents and in nonhuman primates.

## 2 Results

### 2.1 Overview of the Neural Shape Metrics Framework

Consider a set of neural recordings collected across *K* biological systems—for example, *K* different animals, *K* behavioral sessions from the same animal, or recordings from *K* different brain regions (*Fig*. 1a-c). The activity of each neural system is recorded at *M* measurement points, corresponding to different experimental conditions as well as different time points. For example, neural responses could be measured in response to *C* conditions (e.g. distinct sensory stimuli or trained motor actions) over *T* time points per condition. This would result in a total of *M* = *ST* measurements.

**Figure 1:**
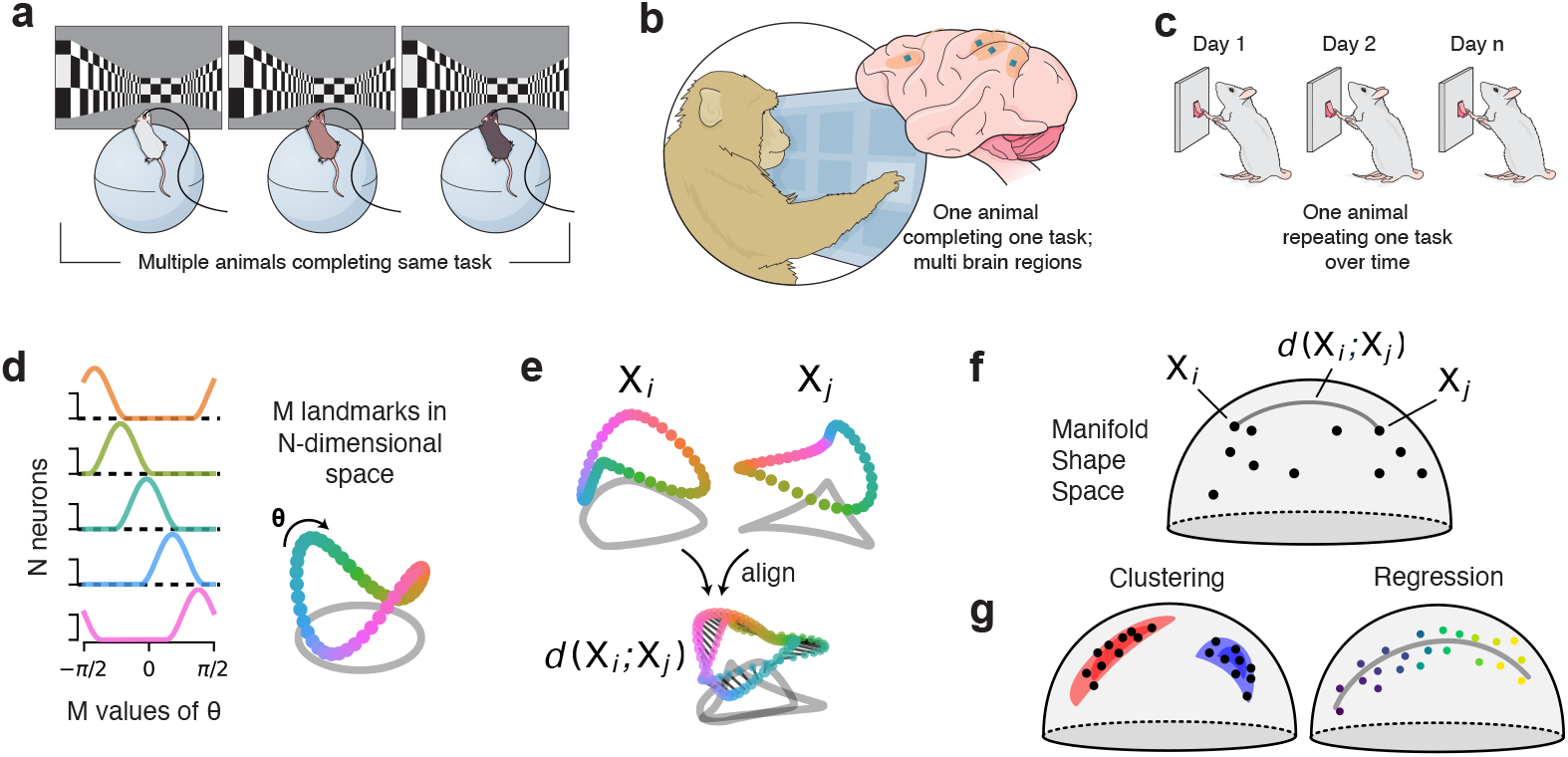
Shape metrics workflow. **a-c**. Shape metrics can be leveraged to compare representations across animals (a), regions (b) or different time periods from the same animal (c). **d**. Example population activity measured across *M* conditions and *N* neurons in an experiment parameterized by a periodic 1D variable, *θ* (e.g. head direction). Smooth tuning curves, *left*, can be viewed as a smooth manifold, *right*, in *N* -dimensional firing rate space. Color code denotes values of *θ*. **e**. The goal of our framework is to compute a distance function *d*(*X*_*i*_, *X* _*j*_), which we define as the dissimilarity between two manifolds *X*_*i*_, *X* _*j*_ after aliment. **f**. The set of possible manifolds and the distance function together define a *metric space*. **g**. The proprieties of these metric spaces allow us to leverage existing methods, such as clustering and regression with rigorous theoretical guarantees.

Our high-level goal is to characterize the extent to which these *K* neural systems are similar or different from each other across these measurement points. To simplify the exposition, we will assume that each neural population is composed of *N* neurons. This assumption is often violated. However, all the methods we outline can be extended to the case of unequal neural population sizes (see *Methods*).

We will focus on comparing trial-averaged neural responses (see *Discussion* for extensions that consider single-trial statistics). In this setting, the data collected from *K* neural systems are represented by a set of *N* × *M* matrices, ***X*** _1_, ***X*** _2_, …, ***X*** _*K*_. Each row corresponds to a neuron’s tuning curve the *M* measurement points (task conditions and time points). Each column corresponds to an *N* -dimensional population response to one of the experimental measurement. *Figure* 1d illustrates an example where the measurements span a 1-dimensional space, resulting in a 1-dimensional, smooth manifold structure within the neural firing rate space.

Our first objective is to define a family of distance functions *d*(***X*** _*i*_, ***X*** _*j*_) that reasonably quantify the dissimilarity between two neural systems. The principal challenge is that the neurons are randomly sampled and labeled in each system and so cannot be immediately compared. This can be interpreted geometrically as two neural manifolds being randomly oriented with respect to each other in firing rate space (*Fig*. 1e, top). Although neural dimensions are not matched between systems, the *M* measurements are shared (coloring in *Fig*. 1d-e). Thus, we can consider *aligning* the two manifolds through some transformation and then computing some measure of distance (*Fig*. 1e, bottom).

We will argue that it is useful to focus on distances which satisfy the following three axiomatic properties:

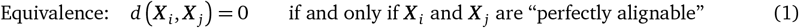

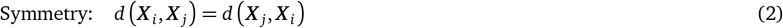

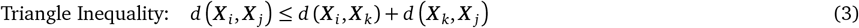

where ***X*** _*i*_ and ***X*** _*j*_ are said to be “perfectly alignable” if the transform class specified by the practitioner is flexible enough to superimpose the two manifolds.

In mathematics, notions of distance that satisfy eqs. (1) to (3) are said to define a *metric space*. Intuitively, metric spaces are useful because they enable us to spatially reason about data. That is, we can envision each of our recordings from *K* neural systems ***X*** _1_, …, ***X*** _*K*_ as points within the space of all possible neural recordings (*Fig*. 1f). That is, in this (potentially high-dimensional) metric space, an entire neural system is represented by a single point, and the distances between points indicate the geometric similarity or dissimilarity of the systems. These properties allow us to leverage methods such as clustering and nearest-neighbor regression (*Fig*. 1g) and to do so with theoretical guarantees (see e.g. Cover and Hart 1967; Dasgupta and Long 2005).

The *Procrustes shape distance* (Schönemann 1966) is a metric that satisfies eqs. (1) to (3), and most of our analyses will leverage this distance as a running example of the shape metrics framework. If we assume that each neuron’s tuning curve has been pre-processed to be mean centered (i.e. rows of ***X*** _*i*_ and ***X*** _*j*_ have mean zero), then the Procrustes distance can be written as:

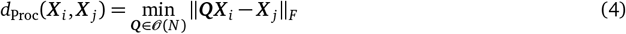

Here, 𝒪(*N*) is the set of *N × N* orthogonal matrices. Intuitively, ***Q*** encodes the optimal rotation and reflection that minimizes the sum of squared residuals between the two neural manifolds. The mean centering step ensures that the rotation and reflection is performed around the manifold’s center of mass.

The Procrustes distance can be thought of as a constrained version of linear regression with an orthogonality constraint—i.e. a constraint such that the weight matrix must only encode a rotation and reflection of the firing rate space. Naïvely dropping this restriction is problematic for constructing metric spaces. Indeed, for a generic *N × N* matrix of regression weights ***W***, the resulting distance score fails to even be symmetric:

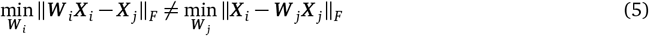

This method of measuring similarity would therefore be sensitive to the order in which systems, such as animals or brain regions, are compared. Thus, simple forms of linear regression do not easily enable comparisons of neural systems within a metric space. Of course, regressions and decoding analyses are still useful tools, and we summarize their complementary advantages in the *Discussion*.

In our experience, the orthogonality constraint inherent to Procrustes distance is often useful to regularize the alignment transformation. Nonetheless, this constraint can be relaxed by computing

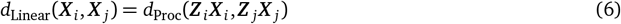

as the distance, where ***Z*** _*i*_ and ***Z*** _*j*_ are *N× N whitening matrices*. This avoids the problem highlighted in eq. (5) while also relaxing the orthogonality constraint inherent to Procrustes distance—i.e. we have that *d*_Linear_(***X*** _*i*_, ***X*** _*j*_) = 0 if any invertible linear transformation brings the two neural manifolds into alignment. The resulting distance is a proper metric and has connections to CCA, as discussed in Williams et al. (2021).

Another, more stringent, metric we will utilize is the *one-to-one matching distance*, which uses a permutation of the *N* neurons to align the responses

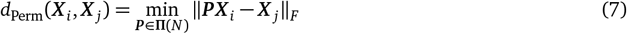

where ***P*** is an *N × N* permutation matrix. Note that the set of permutation matrices, **Π**(*N*), is a subset of all orthogonal matrices, 𝒪 (*N*). Therefore, the Procrustes distance is a less stringent measure of similarity in the sense that *d*_Proc_(***X*** _*i*_, ***X*** _*j*_) ≤ *d*_Perm_(***X*** _*i*_, ***X*** _*j*_). Khosla and Williams (2024) describe how to extend this distance to case where the two populations consist of unequal numbers of neurons.

In this section, we surveyed several metrics that satisfy the desirable properties of eqs. (1) to (3). These belong to an even broader class of distances that are referred to as *generalized shape metrics* (Williams et al. 2021). Additional mathematical details are provided in the *Methods*.

### 2.2 Shape Analysis of 1D Periodic Tuning Curves

We now turn to practical demonstrations. To build intuition, we begin with simulated data from a population of neurons encoding a periodic variable, *θ*. As *θ* is varied, the firing rate traces out a smooth, 1D ring manifold in the high-dimensional firing rate space (fig. 1d). Figure 2a shows tuning curves (*left*) and PCA projections (*right*) from four simulated subjects with *N* = 100 neurons per subject. We simulated similar tuning curves from 36 additional subjects, resulting in a cohort size of *K* = 40.

**Figure 2:**
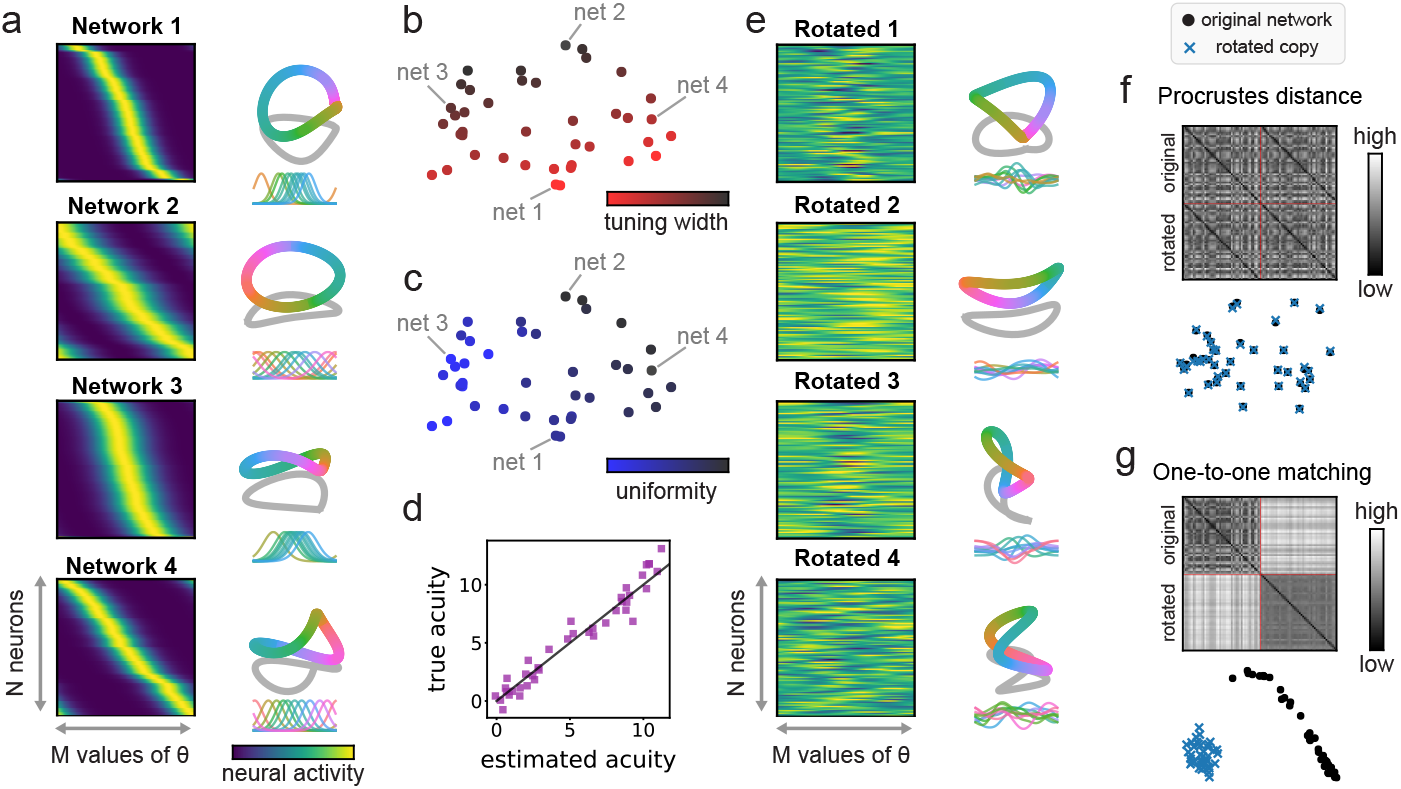
Shape metric analysis of simulated 1D tuning curves. **a**. Left, four example datasets (out of *K* = 40) of *N* = 100 neurons sorted by their preferred stimulus *θ*, with varying tuning width and concentration of tuning around *θ* = 0. Right, first 3 principle components of the same datasets show how tuning curves parameters shape neural manifolds. Inset, five example tuning curves highliting differences in tuning width and uniformity. **b-c**. Embedding of *K × K* matrix dissimilarity matrices in 2D (Methods), shows that the shape metric successfully captured the two forms of across-subject variability, tuning curve width along one dimension (b) and tuning concentration long another (c). **d**. Behavioral performance (true information, calculated using fisher information) is predicted from k = 3 neighbors (*R*^2^ =0.83, estimated information). **e**. Randomly rotated copy of each network in a. **f**. Similar to b, but done for rotated and original networks in e) and a), respectively. Each neural system and its rotated copy lie on top of each other in the low-dimensional space, when using Procrustes distance. **g**. Differences in single-neuron tuning structure in rotated and original networks are are captured by one-to-one matching distance, *d*_Perm_ (see Methods).

In these simulations, we randomly varied two key variables across subjects. First, we varied the average tuning curve width. For example, network 1 has much sharper tuning curves than network 2 in *Figure* 2a. Second, we varied the concentration of tuning around *θ* = 0. In *Figure* 2a, network 3 has many more tuning curves peaked near the center of the horizontal axis, while network 4 has uniformly spaced tuning curves. Non-uniform tiling of tuning curves is seen in neural circuits supporting navigation (e.g., over-representation of reward locations, Hollup et al. 2001) and sensory function (e.g., over-representation of cardinal edges in visual cortex, Li et al. 2003).

We begin by computing a *K × K* matrix of pairwise Procrustes distances, with elements given by ***D***_*i j*_ = *d*_Proc_(***X*** _*i*_, ***X*** _*j*_). We first demonstrate how to use ***D*** to perform dimensionality reduction so that we can visualize the *K* subjects as *K* points in 2D space. We use multidimensional scaling (MDS; Borg and Groenen 2005), which is a well-established technique that takes in a matrix of pairwise distances, ***D***, and returns a list of vectors, ***u***_1_, …, ***u*** _*K*_, such that ∥***u*** _*i*_ − ***u*** _*j*_∥ _2_ ≈***D***_*i j*_ for all pairs of subjects. Some amount of mismatch between the distances is unavoidably introduced by the curvature of the underlying shape space (Robinson 2006; Kendall et al. 2009). However, the distortion is typically small, especially if the embedded vectors, ***u***_1_, …, ***u*** _*K*_, are allowed to occupy a high-dimensional space.

In our simulated example, a 20-dimensional MDS analysis produced achieved a median distortion of ∼3.4% across all pairwise distances (see *Methods*). A 2D PCA projection of this embedding, capturing ∼85% of the variance, is shown in *Figure* 2b-c. From these visualizations, we see the shape metric successfully reveals two forms of across-subject variability that we built into the simulation: subjects are ordered by tuning curve width along one dimension (*Fig*. 2b, top to bottom) and ordered by their concentration around *θ* = 0 along another (*Fig*. 2c, left to right). Thus, variability in tuning curve parameters across subjects are transparently captured within the Procrustes shape metric space.

Now we show how to leverage these distance relationships to make predictions about individual subjects. Suppose, for example, that we are interested in each subject’s ability to perform a sensory discrimination task between two conditions, *θ* = *±ε* where *ε* ≈ 0. A popular way to quantify the ability of a neural representation to support perceptual discrimination is *Fisher Information* (FI), a quantity that is related to manifold geometry and structure of noise (see e.g., Kriegeskorte and Wei 2021). We computed the logarithm of FI per neuron as a proxy for each each subject’s acuity, or theoretical performance on a discrimination task (see *Methods*), and then asked whether we could predict this acuity score for each subject using only their position within the Procrustes metric space. Importantly, we computed the FI acuity score in the asymptotic regime (i.e. assuming an infinite pool of neurons in each network); as described above, our Procrustes distance comparisons are based only on a randomly sampled subset of *N* = 100 neurons. This is meant to simulate a realistic scenario where we only get to measure activity from a small fraction of neurons in each subject.

We found that a *k*-nearest neighbor (kNN) regression model accurately predicted acuity scores in this simulation (*Fig*. 2d, *R*^2^ =0.95, *k* = 3 neighbors). The model works by taking each subject’s *k* nearest neighbors in the Procrustes metric space, and using their average of acuity score as the prediction. Despite its simplicity, this approach can be highly effective, and it has been studied extensively by mathematicians (Cover and Hart 1967). Importantly, practical implementations and theoretical analysis of this model relies on the metric space axioms stated in eqs. (1) to (3).

In our simulation, the kNN performance is improved by increasing the total number of networks, *K*, as well as the number of recorded neurons per network, *N*. Intuitively, increasing *K* improves the model by creating a denser sampling of the Procrustes metric space, while increasing *N* improves our estimate of each subject’s position in the Procrustes metric space.

### 2.3 Rotation vs. Permutation Invariant Shape Metrics

In the previous section, we explored variability across subjects using a rotation-invariant shape metric (Procrustes distance). However, this represents only one choice from the broader family of generalized shape metrics. To demonstrate this, we generate a randomly rotated copy of each network. This results in a neural representations with the same geometry but completely rearranged tuning (*Fig*. 2e). By construction, each rotated copy is indistinguishable from its original counterpart in terms of Procrustes distance. Thus, when we visualize all of the simulated subjects within Procrustes shape space, we see that each neural system and its rotated copy lie on top of each other (*Fig*. 2f).

The Procrustes distance does not necessarily detect differences in the structure of the tuning curve, since many shapes of the tuning curve can give rise to the same geometry at the population level (Kriegeskorte and Wei 2021). However, other shape metrics may be used to quantify and detect these differences. To show this, we use the metric in eq. (7) that is invariant to permutations of the neuron labels but sensitive to rotations (see *Methods*). When we repeat the procedure above with this new distance measure, we see that the rotated copies are positioned far away from their original counterparts (*Fig*. 2g). These two clusters (original vs. rotated networks) are also readily revealed by hierarchical clustering applied to the *K × K* distance matrix.

### 2.4 Animal-to-animal variability in head direction coding

Turning our focus to experimental data, we show how shape metrics can illuminate animal-to-animal variability in two recently published datasets of head direction cells (Duszkiewicz et al. 2024). In both datasets, spiking activity in the mouse postsubiculum (PoSub) was collected using silicone probes during open-field exploration while position and heading direction were captured by an overhead video camera (for details, see Duszkiewicz et al. 2024). The first dataset consisted of *K* = 31 sessions / subjects and a total of 2691 neurons. We first collected single-unit tuning curves for heading across all sessions and sorted them according to peak activity. Preferred heading directions were roughly uniformly distributed (*Fig*. 3a) and, unsurprisingly, applying *PCA* to this aggregated dataset revealed a high-dimensional ring structure (*Fig*. 3a, bottom), akin to the simulated dataset we analyzed above.

**Figure 3:**
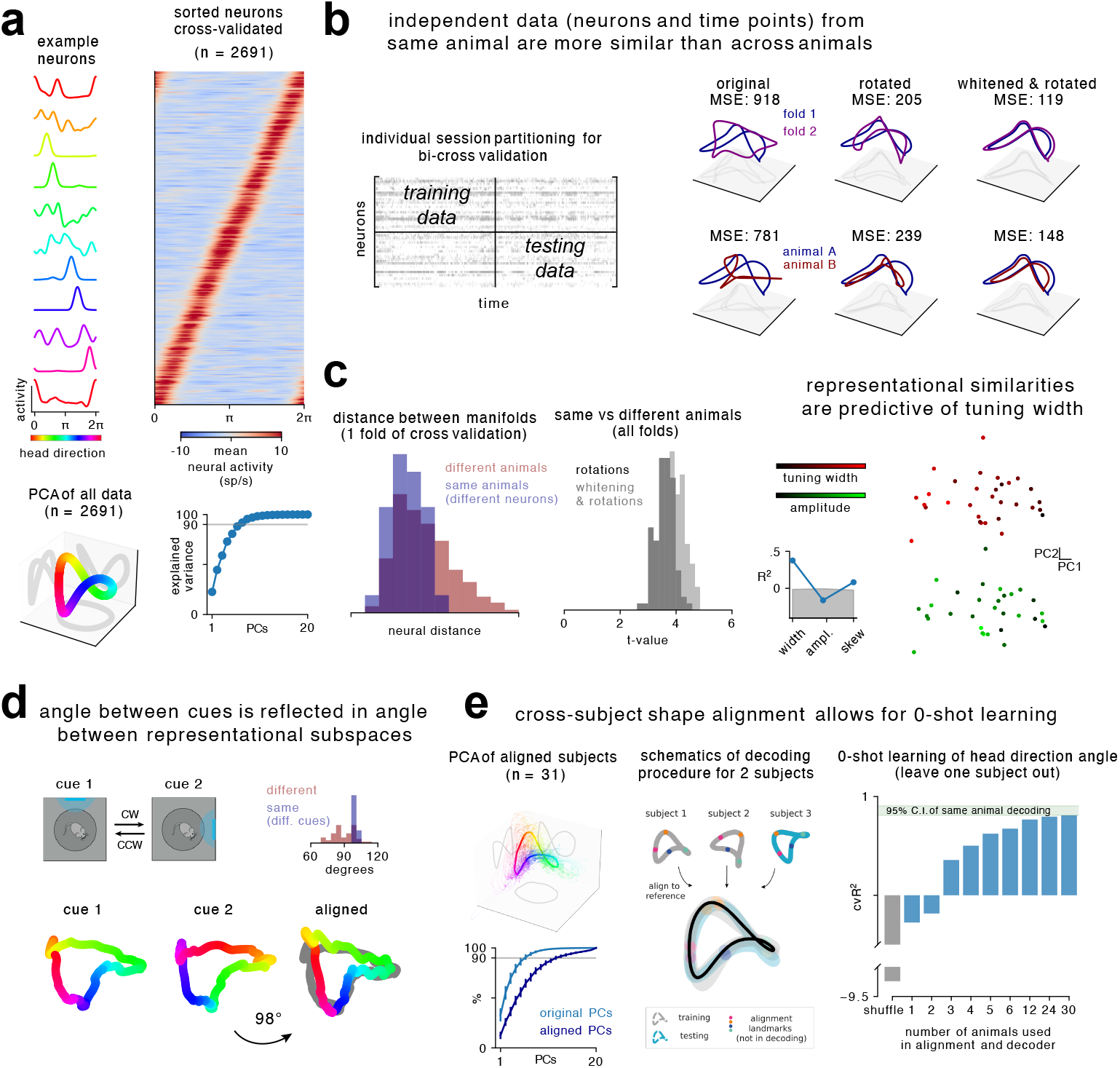
Shape metric analysis of head direction coding. **a**. Left, example neurons. Right, all neurons sorted (cross-validated across time) by their preferred direction angle. Bottom, top three principal components of all neurons reveal a high-dimensional ring. **b**. Left, illustration of the cross-validation approach used for this dataset. Right, independent folds of animal A (blue and purple, top) have smaller mean squared-error (MSE) distance after alignment than a similar analyses between animal A and B (blue and red, below). **c**. Same analyses in b), repeated for all animals and 1 individual split. Distance between two different splits of the same animal are significantly smaller than across different animals (t=-3.13, p=0.0017). Middle, the former replicated on n=100 different splits. Whitening the data before alignment captures individual differences slightly better (p=0.05). MDS of the dissimilarity matrix suggests that tuning width drives across-animals differences (red gradient vs green gradient). Inset, decoding *R*^2^ (using bi-cross validation in b) of tuning width from this space is significantly above chance (95% C.I. of shuffle decoder in gray). **d**. Analyses on a separated dataset of n=6 animals shows that a rotation in the cue space (90°) is reflected in a consistent angle rotation in the neural space (98°). **e**. Left, aligning all subjects (n=31) to a common reference (Methods) reveals a high-dimensional ring. Bottom, the first principal components of the aligned space (dark blue) explains a substantial amount of variance of the original space (light blue). Right, illustration of a 0-shot learning decoding procedure form 2 aligned animals. As we aligned a larger number of subjects, the decoder approaches the performance of the decoder trained and tested on independent folds of the same animal.

To our knowledge, it is unknown whether there are systematic animal-to-animal differences in the statistics of head direction tuning curves in PoSub. To investigate, we measured the Procrustes distance between two disjoint partitions of the data along neuron and time dimensions (see illustration in *Fig*. 3b, left; *Methods*). On average, the distance between partitions of the same animal was smaller than the distance between partitions taken from different animals (*Fig*. 3b-c). Whitening the data before alignment, as in eq. (6), slightly improved the capture of individual differences (*Fig*. 3c, middle).

What features of the neural tuning curves account for these individual differences in ring shape? To perform an exploratory analysis, we applied MDS followed by PCA to visualize each session in a low-dimensional space To investigate, we first used unsupervised methods to perform an exploratory analysis. Akin to our analysis of the simulated data above (*Fig*. 2b-c), we used MDS followed by PCA to visualize each subject within a 2D space. MDS introduced a median distortion ∼2% and PCA captured ∼64% of variance in the MDS embedding. Coloring each session by the average width and average amplitude of the head direction tuning curves suggested that the former, and not the latter, was a principal driver of inter-animal differences (*Fig*. 3c, right). To confirm this finding in our exploratory analysis, we turned to supervised methods. The decoding the tuning width from this space was significantly above zero (bi-cross validated as in *Fig*. 3b), demonstrating that indeed tuning width is driving the inter-individual differences captured by the shape metrics (*Fig*. 3c, inset).

We found similar signatures of individual differences in a second, smaller dataset (*K* = 6 animals). In this experiment, open exploration was performed in boxes with a salient landmark which was rotated by 90 degrees during the course of the recording (cue 1 vs cue 2, *Fig*. 3d). We find that the optimal rotation identified by Procrustes analysis in neural firing rate space was roughly matched in magnitude to the cue rotation (*Fig*. 3d).

The analyses above show how the Procruste distance can be used to detect individual differences. We now show that a similar analysis can identify common manifold structure across subjects and leverage this to decode head direction in one heldout subject from training data recorded in other subjects. We started by using rotation and reflections to align all subjects to a common reference—a procedure known as *generalized Procrustes analysis* (Gower 1975) or *hyperalignment* (Haxby et al. 2020). This revealed a high-dimensional ring structure (*Fig*. 3e), similar to performing PCA on all data (*Fig*. 3a) but now confirming that this structure is shared by all subjects and not just present in a subset of them. In fact, the first principal components of this aligned space explained a substantial amount of variance of the original space of each animal (*Fig*. 3e, left). Because all animals are aligned to a common shape (in this case, a ring), we could train a decoder on a subset of animals and predict head direction angles on left-out subjects, essentially performing zero-shot learning (*Fig*. 3e, right).

In sum, all subjects encode the head direction angle in a ring, but each of these rings have idiosyncratic distortions in higher dimensions. Both common structures and individual differences can be analyzed using shape metrics.

### 2.5 Measuring representational similarity across numerous brain regions

Next, we investigate how shape metrics can be used to reveal detailed functional relationships across brain regions. To demonstrate generality of this approach, we performed two separate analyses of data from mice and nonhuman primates. In both cases, we interrogate the conventional understanding that population-level representations morph along an anatomical hierarchy starting at early visual areas (e.g. primary visual cortex) and culminating at higher-order areas (e.g. parietal and prefrontal cortices).

First, we studied population-level activity across *K* = 55 fine-grained subregions of mouse cortex and thalamus in data publicly released by the Allen Brain Observatory (Siegle et al. 2021). We focused on extracellular electrical recordings (Neuropixels probes) performed while mice viewed a naturalistic movie (Touch of Evil by Welles, O.;, *Fig*. 4a). Spike counts attributed to isolated units were trial-averaged, normalized, and pooled across subjects before categorizing them into *K* = 55 populations based on anatomical location defined by the Allen Mouse Brain Common Coordinate Framework and Allen Mouse Brain Atlas. As done above, we computed a *K × K* matrix of pairwise Procrustes shape distances across these subregions (*Fig*. 4b), and visualized the result using MDS and PCA (*Fig*. 4c). MDS introduced a median distortion of ∼2.3% in the pairwise distances. The top two principal components captured only ∼22.5% of the variance in the MDS embedding, but nonetheless revealed interpretable structure. In particular, coloring each point within this 2D space according to the anatomical atlas conventions (*Fig*. 4a, inset) reveals that the subregions cluster together according to anatomy (e.g. the separation of thalamic and cortical areas in pink vs dark green in *Fig*. 4c). The metric properties of shape space enable us to rigorously quantify this intuition: applying hierarchical clustering, an unsupervised learning method for clustering data points, to the dissimilarity matrix we find that areas with similar anatomical position are clustered together (*Fig*. 4b).

**Figure 4:**
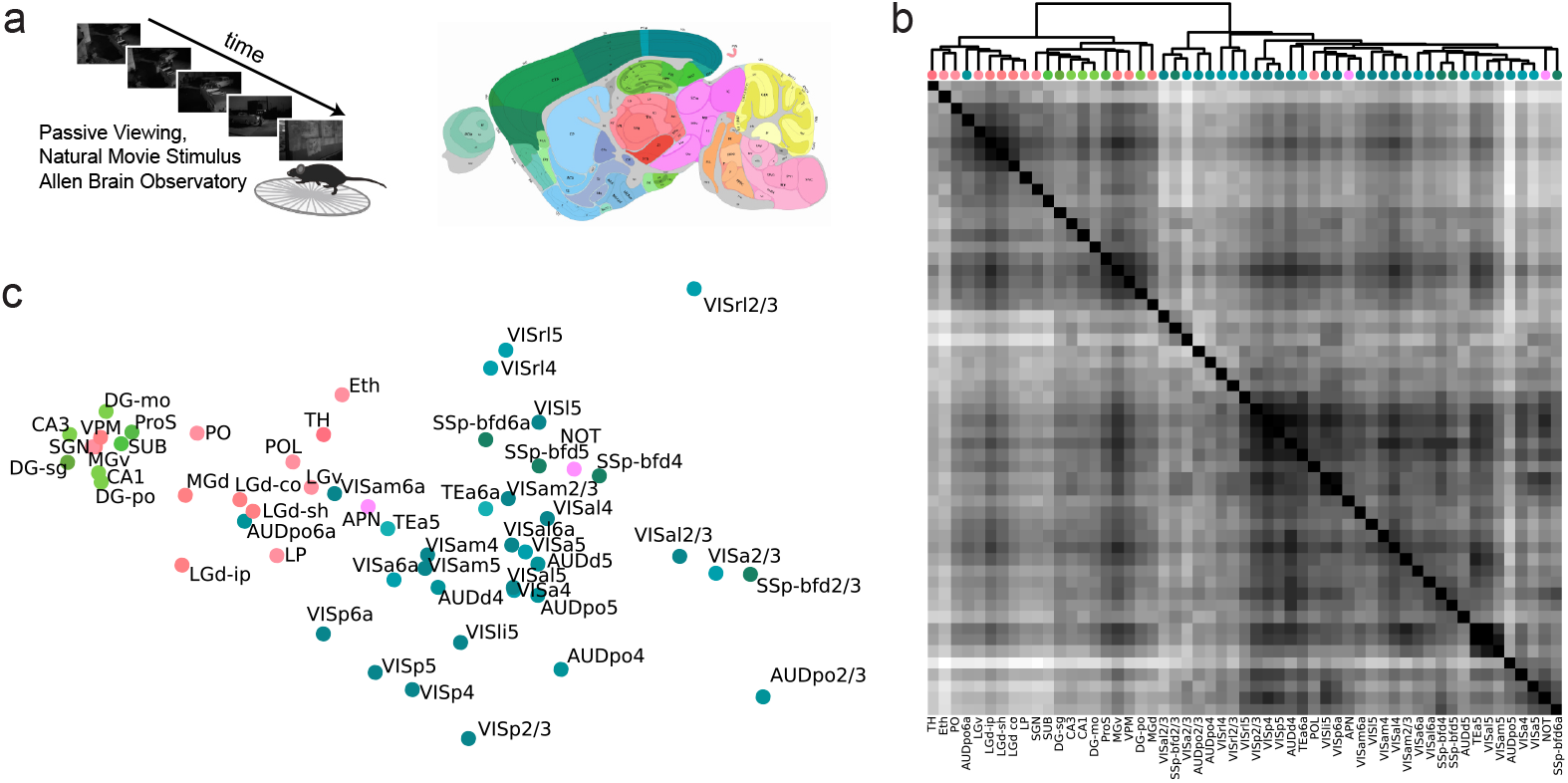
Comparison of visual representations across brain regions in mice. **a**. Illustration of the Allen Brain Observatory dataset, consisting of K = 56 fine-grained subregions of mouse cortex and thalamus (Right) in response to a naturalistic movie (Left). **b**. K x K matrix of pairwise shape distances across all subregions. The matrix is further sorted by clusters found using hierarchical clustering (Methods) **c**. Visualization of b), using MDS and PCA shows subregions cluster together according to anatomy.

In a second analysis, we examined data from two nonhuman primate subjects performing a complex decision-making task (*Fig*. 5a) while recording from *K* = 7 regions, published in Siegel et al. (2015). Calculating pairwise shape distances between all areas and independent partitions of the dataset for each area (*Methods, Fig*. 5b) revealed several interesting features that are noticeable when the dissimilarity matrix visualized in 2D (*Fig*. 5c). Namely, we found that regions were organized by anatomical hierarchy, that independent partitions (connected in *Fig*. 5c) were closer to each other than to other regions and that similar areas of different subjects (different shades of the same color) were more similar to each other than to other regions. We quantified these interesting features in the following way. First, the anatomical location (Fig 5d,e) and the subject (Fig 5e, right) of individual brain regions could be predicted with threshold-based ordinal logistic regression trained and tested in independent partitions of the dissimilarity matrix (*Methods, Fig*. 5d). Second, independent partitions of the same region are more similar to each other than partitions of different regions (*Fig*. 5f, left). Finally, the same areas (e.g. PFC R and PFC P) of different animals are more similar than different areas (*Fig*. 5f, right).

**Figure 5:**
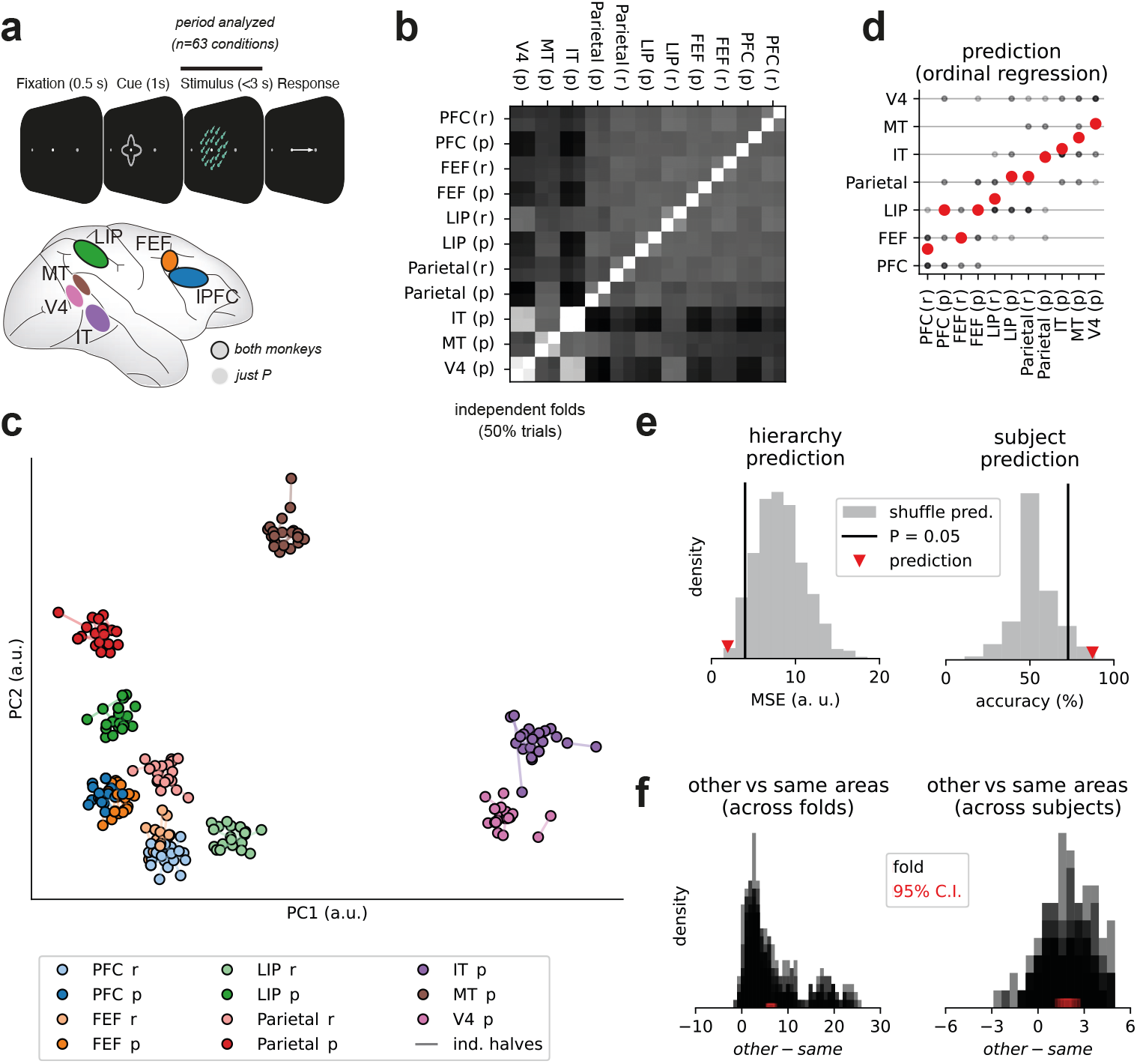
Deterministic shape metrics can be used to quantify encoding geometry differences across regions. **a**. Description of a multi-region dataset analyzed with deterministic metric. **b**. Matrix of dissimilarities between all areas of each subject and two independent partitions. **c**. PCA visualization of B. Dark colors are from subject P and light colors from subject R. Different dots (n=10) are different repetitions of the analyses, aligned with Procrustes analyses for visualization. Independent partitions are connected by a colored line. **d**. Areas can be predicted using ordinal regression (cross-validated). Black circles mark predictions of each n=10 partitions, and red circles the average prediction. **e**. Both hierarchy and subject identity can be predicted above chance. **f**. Areas are more similar to each other than to other random areas. Left, distance calculated between independent partitions of the same area. Right, distance calculated across subjects.

In sum, by using shape metrics to compare neural representations across brain regions, we have recovered anatomical position in an unsupervised fashion in both mice and monkeys. Moreover, by exploiting the metric properties of shape space we could perform both supervised and unsupervised learning methods on that space (i.e. linear regression and clustering, respectively) and rigorously quantify hypotheses developed through visual inspection (e.g. region organization by anatomical hierarchy in PCA space was quantified using regression or clustering in the full space).

### 2.6 Relating individual differences in behavior to differences in neural representations

As a final example, we used shape metric analysis to investigate whether differences in behavioral performance are reflected in differences in neural population activity. To do this, we turned to public data released by the International Brain Lab (International Brain Laboratory et al. 2024). Two features of this dataset make it an ideal candidate for this analysis. First, it consists of neural and behavioral measurements from a large cohort of animals performing the same binary choice task involving contrast discrimination (*Fig*. 6a). Second, a large number of neurons were measured in each animal spanning the same brain regions including secondary visual areas, hippocampus, and thalamus. These regions have previously been shown to encode all task variables (International Brain Laboratory et al. 2024).

**Figure 6:**
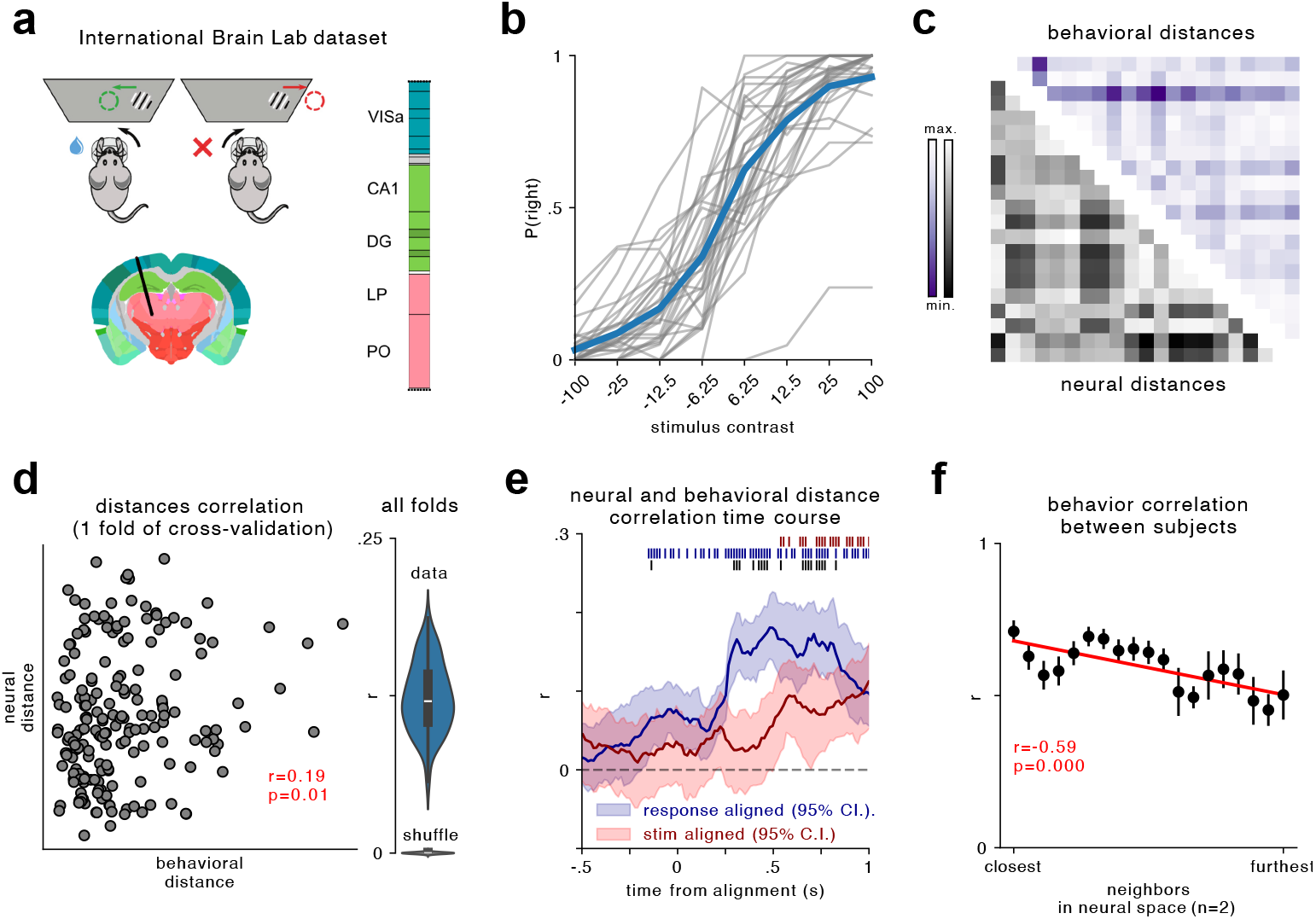
**a**. IBL dataset schematics. **b**. Psychometric curves all animals (n=20) in gray and grand average in blue. **c**. Matrix of neural and behavioral dissimilarities between all animals. Note that in all the analyses, independent data was used to compute behavioral and neural distances. Here shown for one data split. **d**. Left, correlation between behavioral and neural distances of one fold. Right, correlation values of n=100 repetitions of the data split and similar analyses for shuffled data. Both done using neural data around the response ([0-0.4 s]). **e**. The correlation between neural and behavioral distances throughout the trial and aligned to stimulus or response. Red, blue and black ticks on the top mark the time-points where stimulus, response or stimulus vs response was significantly different than 0 (bootstrap t-test), respectively. **f**. Alignment between neural representations allows for 0-shot learning of psychometric curves using the 2-closest neighbors. Behavioral correlation between the first (2 closest neighbors) and last point (2 furthest neighbors) is significantly different (*t* = 2.2, *p* = 0.037).

The original dataset contains 78 mice; however, we excluded mice with fewer than 100 neurons or 20 behavioral trials per condition. This limited our analysis to *K* = 20 subjects. To quantify the behavioral profiles on the task, we computed psychometric curves as a function of stimulus contrast. These curves display considerable variability across subjects (*Fig*. 6b), prompting us to ask whether there is a neural signature of this variability.

We started by computing matrices of pairwise distances across subjects quantifying neural variability (Procrustes metric, eq. (4)) and behavioral variability (mean absolute distance between psychometric functions). Importantly, we used independent trials to calculate the Procrustes and behavioral distances between animals to avoid spurious correlations due to shared noise. A correlation analysis between the behavioral and neural distances of the particular partition shown in (*Fig*. 6c) revealed a significant relationship between these two measures, calculated on independent data (*Fig*. 6d). Repeating this analysis 100 times with different data partitions confirmed that the correlation values were consistently higher than the correlations between shuffled subjects (*Fig*. 6d). This analysis confirmed that behavioral strategies are reflected in the encoding geometry. To pinpoint what drove this relationship, we performed a time-resolved analysis.

Specifically, we examined the correlation between neural and behavioral distances, as above, but throughout the trial. Moreover, we repeated this analysis aligning the data with stimulus or response times. Our results showed significant differences at various time points, marked by red, blue, and black ticks for stimulus, response, or stimulus vs. response, respectively (*Fig*. 6e). As expected, above-chance correlations were found before the stimulus presentation. Instead, this correlation built up in course of the trial. Interestingly, we found the strongest correlation when focusing on response-aligned data, including a week but significant correlation before the response was taken. This analysis suggested that the relationship between behavioral strategies and encoding geometry was dominated by response-related changes in the encoding geometry.

Finally, we show once again (see also Figs. 2 and 3) how aligning neural representations allows for zero-shot learning of psychometric curves using the two closest neighbors. Specifically, for every subject in the dataset we computed the behavioral correlation between this subject and the two nearest neighbor subjects in the neural shape space (closest, in *Fig*. 6f). We then repeated this analysis, but now predicting each subject behavior using progressively distant subjects in neural shape space. We found that the behavioral correlation between the first (two closest neighbors) and the last (two furthest neighbors) was significantly different (*t* = 2.2, *p* = 0.037). Furthermore, we found a negative correlation between distance in neural shape space and behavioral distance (*Fig*. 6f), showing that the behavioral correlation decreases smoothly for increasingly distant animals. All together, these analyses show how different behavioral strategies are reflected in different neural geometries, as quantified by shape metrics.

## 3 Discussion

Optical and electrophysiological neural recordings have exponentially increased in scale over the past several decades (Stevenson and Kording 2011; Urai et al. 2022). These methods are now maturing into plug-and-play technologies that are commercially available at increasingly affordable rates, raising the possibility of scaling up neural datasets in fundamentally new ways. In particular, population-level neural recordings can now be obtained across many subjects, brain regions, and repeated behavioral sessions, but summarizing these multi-session datasets is a formidable challenge. While prior works have introduced relevant methods (see Klabunde et al. 2023, for a review), future advances will require us to further develop and refine these approaches. Here, we demonstrated the potential of a broad and flexible framework based on shape metrics.

Systematic comparisons of neural population recordings will require a variety of approaches including both unsupervised methods for exploratory analysis (e.g. dimensionality reduction and clustering) as well as supervised methods for testing targeted hypotheses (e.g. regression and classification models). We showed that a number of these analyses are enabled by establishing notions of distance that are symmetric and satisfy the triangle inequality (i.e. define a metric space). For example, we used clustering in Procrustes shape space to identify brain regions with similar neural population activity, and found that these clusters were in line with anatomical knowledge (Fig. 4,5). Such analyses may prove useful in the context of ongoing debates about the extent of functional modularity within the brain (see e.g. Ostojic and Fusi 2024).

We also demonstrated that shape metrics can be leveraged to fit supervised models. For example, a neural system’s position within Procrustes shape space can be used to predict single-neuron proprieties (Fig. 3), anatomical location (Fig. 5) and psychometric performance (Fig. 6). The final example relating individual differences in neural population dynamics to differences in behavior seems to be a particularly compelling use case that few prior works have attempted. An important exception is recent work by Fascianelli et al. (2024), which performed a detailed analysis of two nonhuman primate subjects and found correlations between their behavioral differences and neural geometries. Shape metrics provide a rigorous framework to scale up these investigations to larger experimental cohorts (e.g. across 20 mice in Fig. 6).

Shape metrics are not the only family of distance functions that satisfy symmetry and the triangle inequality. Slight modifications to CKA scores and CCA-based similarity scores result in proper metric spaces (Williams et al. 2021). Distance-based RSA scores can also be turned in proper metrics, as can be seen from recent work showing equivalence between RSA and CKA (Williams 2024). Despite these alternatives, our work is one of the first to emphasize and demonstrate the advantages of using proper metric spaces on neurobiological data.

Linear predictivity scores are another framework for comparing neural systems. They are particularly popular for quantifying similarity between artificial and biological networks (e.g. Conwell et al. 2022). One can view shape metrics as a framework for incorporating constraints into predictivity scores in order to turn them into proper metric spaces. Indeed, raw predictivity scores can be arbitrarily asymmetric—it is easy to construct a synthetic example such that predicting neural system *A* from *B* yields *R*^2^ ≈ 1 while predicting neural system *B* from *A* yields *R*^2^≈ 0 (see *Methods*). This asymmetry is not strictly a defect of this method. In fact, it is desirable in circumstances where one only cares about predicting *A* from *B* (and not *B* from *A*). This might be the case, for example, when artificial deep networks are used to predict biological responses for the purpose of running *in silico* experiments. Here, we have focused on a different problem of performing comparative analyses among many biological systems. Since there is often no *a priori* asymmetry between the systems we would like to collectively evaluate, it is natural to compare them within a proper metric space.

For simplicity, we utilized the Procrustes distance as a running example throughout most of our analyses. However, this is just one of a broader class of possible shape metrics. Our practical experience suggests that the Procrustes distance is reasonable default choice—it is resistant to overfitting and has no hyperparameters that need to be tuned—but practitioners should consider a broader set of distance functions depending on their analysis goals. For example, the Procrustes distance is not sensitive to differences in the tuning structure of individual neurons, as we demonstrated in Fig. 2e-g. A more stringent distance measure is given by the one-to-one matching distance, eq. (7), and its generalization, the *soft matching distance* (Khosla and Williams 2024). These alternatives can be used to quantify the extent to which neural systems represent information in arbitrarily rotated coordinate systems or in terms of a privileged neural basis (Khosla et al. 2024).

While the flexibility of the shape metrics framework can be advantageous, it also highlights a challenge. Shape metrics represent only a fraction of a broader array of methodologies—a recent review by Klabunde et al. (2023) cataloged over thirty different representational similarity measures from the machine learning literature. Many of these could, in principle, be applied to neurobiological data. How should practitioners choose among the large collection of methods that quantify similarity in neural population codes?

While there is no simple answer to this question, shape metrics provides a unifying conceptual framework that can help address the underlying challenge. By defining similarity in terms of explicit geometric alignment transformations (e.g. rotations or permutations of neural firing rate space), this framework forces practitioners to be explicit with their modeling assumptions. Furthermore, it enables us to reason about and compare different analysis outcomes. For example, we know that the Procrustes distance between two systems will *always* be less than or equal to the one-to-one matching distance because the former allows for a more permissive set of alignment transforms. Neuroscientists can use these relationships among shape metrics to interpret and compare different forms of analysis.

By grounding comparisons of neural systems in explicit alignment transformations, shape metrics are related to a number of previous analyses. For example, Chen et al. (2021) analyzed hippocampal population activity and found that they could use rotational alignments to predict neural responses in one condition (e.g. right turns on a T-maze) using cross-subject alignments on a different condition (e.g. left turns). Similar cross-condition decoding analyses have been applied elsewhere (Saez et al. 2015; Bernardi et al. 2020; Barbosa et al. 2023). Here we showed that such rotational alignments can be used to define a metric space, and therefore used to rigorously analyze larger collections of neural systems (not just pairs of subjects or conditions). However, there is a clear relation and potential synergy between these analytic approaches (see e.g. Fig 3e). In contrast, it is more difficult to draw direct connections to alternative representational similarity measures, like CKA and RSA, which are not defined in terms of explicit geometric transformations (but see Harvey et al. 2024, for a relation between Procrustes distance and RSA-style methods).

The potential of the shape metrics framework to study neural geometry was not fully explored here. We have focused on analyzing trial-averaged neural responses, but the scale and orientation of trial-to-trial noise is important to theories of neural coding (Moreno-Bote et al. 2014). Furthermore, there is substantial interest in moving beyond geometry to understand neural circuits as dynamical systems (Vyas et al. 2020). The framework we’ve outlined can be extended to address these issues (Duong et al. 2023; Nejatbakhsh et al. 2024; Ostrow et al. 2024). However, these extensions remain a relatively active area of research.

Overall, we’ve aimed to show that the simplest forms of shape metric analysis are viable off-the-shelf tools to perform comparative systems-level analysis of neural data. We focused primarily on Procrustes shape distance, which has no tunable hyperparameters and is both analytically and computationally tractable. Leveraging the metric space properties of this distance, we revealed several novel and non-trivial insights including across-subject differences in the population geometry of head direction codes, clustering and functional modularity of representations across brain regions, and a correlation between neural population geometry and behavioral performance. As recording technologies continue to advance, generating increasingly large and complex neural datasets, shape metrics have the potential to become a fundamental tool for neuroscience, enabling researchers to systematically analyze large collections of neural systems at the level of population activity.

## 4 Methods

All experimental data analyzed in this study was previously collected and described in several publications (Siegel et al. 2015; Siegle et al. 2021; Duszkiewicz et al. 2024; International Brain Laboratory et al. 2024).

### 4.1 Datasets and Analyses

#### 4.1.1 Simulated Dataset (Fig. 2)

We randomly generated *K* = 40 sets of tuning curves (corresponding to different “networks” or “subjects”). Each simulated network consisted of *N* = 100 neurons with tuning curves given by a 2-parameter family of functions:

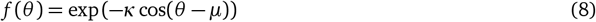

Intuitively, *µ* ∈ [−π, π) describes the peak of the tuning curve and κ ≥ 0 inversely controls the width of the tuning. Under this model, the peak firing rate of each neuron is equal to one. Note that eq. (8) is proportional to the density of a von-mises distribution, for which κ is called the concentration parameter and *µ* is called the location parameter.

To generate the dataset, we specify a value of *κ* and a distribution over *µ* for each subject and sample *N* = 100 random tuning curves. The concentration parameter for each subject, κ, was sampled uniformly between 1 and 4; all tuning curves from each subject were assigned the same κ parameter. The location parameter for each tuning curve, *µ*, was sampled from a von-mises distribution with mean zero and a concentration parameter *γ* that was drawn uniformly between 0 and 3 for each subject. Intuitively, a simulated network will roughly have uniformly spaced tuning curves when the concentration parameter controlling the distribution of *µ* is close to 0. Conversely, a simulated network will have tuning curves that are “bunched” around *θ* = 0 when this concentration parameter is close to 3.

We computed the visual acuity of each subject by taking the logarithm of the expected Fisher Information (FI) across neurons evaluated at *θ* = 0. Specifically, under the assumption of isotropic Gaussian noise with variance *σ*^2^, the expected FI is given by

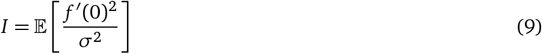

where the expected value is taken over randomly sampled tuning curves, *f*. Note that in our simulated data we only get to observe *N* = 100 tuning curves to compute the Procrustes distances between network pairs. This simulates a realistic scenario where we only get to experimentally record a small fraction of the neural circuit underlying behavioral performance. Nonetheless, if *N* is large enough and the shape of the neural representation is smooth/low-dimensional, we can expect the empirical Procrustes distance to do a reasonable job of capturing the “true” geometry of the system.

Applying some basic algebraic manipulations to eq. (9), we obtain

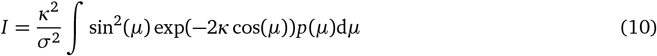

where *p*(*µ*) is the density of a von-mises distribution with mean *θ* = 0 and concentration parameter *γ*. Although some further simplifications are possible, the form of the resulting integral does not appear to have a simple analytic solution. Thus, we numerically computed it to high precision to obtain log *I*, the simulated “acuity” score for each subject. We set *σ*^2^ = 1 in this calculation; this choice is of little consequence since it only introduces an overall scaling into the simulation.

#### 4.1.2 Head direction coding in mice (Fig. 3)

We analyzed two datasets made public by Duszkiewicz et al. (2024). Spiking activity from cells in the mouse postsubiculum (PoSub) was collected using silicone probes during open-field exploration. The first dataset consisted of 31 sessions/subjects and a total of 2691 neurons. The second dataset is similar to the first one, but included a salient landmark that was rotated 90° back and forth between every 200s for a total of 16 rotations.

In both datasets, spikes were counted in bins of 1° of the head direction and then bins were smoothed with 12° s.d. Gaussian filters, as done in the original publication (Duszkiewicz et al. 2024). Each partition for cross-validation (see below) was then denoised by keeping only the first 20 principal components (see Fig. 3a).

To perform leave-one-subject out decoding, we aligned all subjects to a common reference. The common reference was chosen randomly and did not change the results. We aligned each subject manifold using n=20 randomly chosen, but equally spaced, head direction bins. Importantly, we did not include these angles nor *±*6 bins around them in the training set for head-direction decoding.

#### 4.1.3 Allen brain observatory dataset (Fig. 4)

We analyzed “natural movie three” stimulus set of the Neuropixels—Visual Coding dataset in the Allen Brain Observatory (Siegle et al. 2021). Data were obtained and analyzed through the AllenSDK python package. We pooled data from 32 recording sessions, which is the total number of sessions employing the *brain observatory 1*.*1* stimulus set. Spike counts were binned at the video presentation frame rate (30^−1^ second time bins), averaged across ten trials, and smoothed by convolving with a Gaussian kernel (standard deviation of 20 time bins). This resulted in *M* = 1200 time bins of activity measurements for each neuron. Neurons were then pooled across sessions and separated into subregions using the most fine-grained anatomical regions defined by the Allen Brain Atlas. We excluded brain regions with fewer than 50 neurons and additionally excluded subregions with the acronyms `{MB, alv, ccs, dhc, fp, or}` which represent either white matter or midbrain neurons that were not assignable to any specific nucleus. This resulted in a total of *K* = 55 brain regions. To denoise these datasets and speed up calculations, we projected each dataset onto the top *N* = 50 principal components before analysis.

#### 4.1.4 Nonhuman primate visually guided decision-making task (Fig. 5)

Acute recordings from 7 regions of the monkey cortex were performed using single tungsten/paralyene electrodes (Siegel et al. 2015). Monkeys were trained to performed a context-dependent decision-making task. The stimulus consisted of dots moving to a random direction and of a random color.

We computed conditioned-averaged responses (n=64 conditions, 4 colors x 4 directions x 4 contextual cues; Siegel et al. 2015) from *N* = 7542 multiunits recorded from 4 brain regions from subject P, and 7 regions from subject R (Fig. 5A). For each condition, we averaged two independent sets of trials separately (folds, in Fig. 5). We repeated this processed 10 times, leading to 10 datasets consisting of 2 independent folds of 11 brain regions. Finally, to avoid across-area differences to be dominated by different dataset sizes, we subsampled each dataset to n=80 neurons and reduced their dimensionality to M=5 using PCA, leading to a data tensor of dimensions 2 × 11 × 5 that we analyse using shape metrics. All the analyses were repeated 100 times and we report the averages.

#### 4.1.5 International Brain laboratory dataset (Fig. 6)

We analysed a subset of the IBL dataset published in International Brain Laboratory et al. 2024. Briefly, the IBL labs conducted Neuropixels recordings from the same brain regions in head-fixed mice during a standardized decision-making task, where the mice reported the perceived position of a visual grating. The experiment followed a uniform pipeline across all labs, including standardized methods for surgery, behavioral training, recordings, histology, and data processing. We focused here exclusively on behavior and spike counts recordings.

In addition to the original quality control, we only analyzed subjects from which at least *n* = 100 neurons were recorded and with at least *n* = 20 trials per condition were recorded. Moreover, we only focused on data collected during balanced blocks (i.e. in which the probability of rewarded left and right responses was the same). In total, we analysed data from *n* = 20 subjects. For subjects with more than *n* = 100 neurons, we focused the analyses on the top-100 neurons, sorted by task-related variance.

### 4.2 Shape Metrics and Downstream Analyses

#### 4.2.1 Multidimensional Scaling (MDS)

We use a public implementation of metric MDS from `scikit-learn` (Pedregosa et al. 2011). Briefly, given a *K × K* matrix ***D*** of pairwise distances, MDS uses iterative parameter updates in an attempt to solve the following nonconvex optimization problem

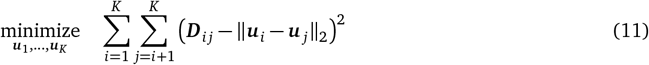

where ***u***_1_, …, ***u***∈ _*K*_ ℝ^*d*^ are vectors within a *d*-dimensional embedding space. Once obtained, these vectors can be used for any downstream analysis (e.g. running PCA to visualize each neural system in a low-dimensional space; see Figures 2b-c, 3c, 4c). However, MDS comes at the cost of introducing distortion into the metric space. We quantify the distortion of distance ***D***_*i j*_ as

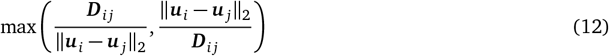

and report the median distortion across all distance values.

## 5 Acknowledgements

This research was supported by the Office of Naval Research (N00014-22-1-2453), Freedom Together Foundation, and the Picower Institute for Learning and Memory.

